# Increased sedoheptulose-1,7-bisphosphatase content in the C_4_ species *Setaria viridis* does not affect photosynthesis

**DOI:** 10.1101/2022.05.09.491242

**Authors:** Maria Ermakova, Patricia E. Lopez-Calcagno, Robert T. Furbank, Christine A. Raines, Susanne von Caemmerer

## Abstract

Sedoheptulose-1,7-bisphosphatase (SBPase) is one of the rate-limiting enzymes of the Calvin cycle, and, in C_3_ plants, increasing the abundance of SBPase is known to provide higher photosynthetic rates and stimulate biomass and yield. C_4_ plants usually have higher photosynthetic rates because they operate a biochemical CO_2_ concentrating mechanism between mesophyll and bundle sheath cells. In the C_4_ system, SBPase and other enzymes of Calvin cycle are localised to the bundle sheath cells. Here we tested what effect increasing abundance of SBPase would have on C_4_ photosynthesis. Using *Setaria viridis*, a model C_4_ plant of NADP-ME subtype, we created transgenic plants with 1.5 to 3.2-times higher SBPase content, compared to wild type plants. Transcripts of the transgene were found predominantly in the bundle sheaths suggesting the correct cellular localisation of the protein. Abundance of RBCL, the large subunit of Rubisco, was not affected in transgenic plants overexpressing SBPase, and neither was relative chlorophyll content or photosynthetic electron transport parameters. We found no correlation between SBPase content in *S. viridis* and saturating rates of CO_2_ assimilation. Moreover, detailed analysis of CO_2_ assimilation rates at different CO_2_ partial pressure, irradiance and leaf temperature, showed no improvement of photosynthesis in plants overexpressing SBPase. We discuss potential implications of these results for understanding the regulation of C_4_ photosynthesis.

## Introduction

Global crop production needs to double by 2050 to meet the projected demands from a rising population, diet shifts, and increasing biofuels consumption (Ray et al., 2013). Increasing photosynthetic capacity of plants was proposed to significantly increase crop yield (Long et al., 2006, Parry et al., 2013, Evans, 2013, Bailey-Serres et al., 2019, Ort et al., 2015, Raines, 2011, Simkin et al., 2019). This has led to research efforts focusing on improving various photosynthetic components in attempt to improve plant productivity (Kromdijk et al., 2016, Ermakova et al., 2021b, López-Calcagno et al., 2019, Lefebvre et al., 2005, South et al., 2019, López-Calcagno et al., 2020).

The Calvin-Benson-Bassham (C_3_) cycle is the primary pathway for CO_2_ fixation in all terrestrial plants. This cycle plays a central role in plant metabolism providing intermediates for starch and sucrose synthesis as well as isoprenoid metabolism and the shikimic acid biosynthesis (Geiger and Servaites, 1994). Manipulation of C_3_-cycle enzymes has led to increases in photosynthetic rates (reviewed by Simkin et al., 2019). In particular, overexpression of sedoheptulose-1,7-bisphosphatase (SBPase) in several C_3_ species has led to increases in photosynthetic rates and increased biomass in the laboratory and the field (Rosenthal et al., 2011, Lefebvre et al., 2005, Driever et al., 2017, Feng et al., 2007, Ding et al., 2016). SBPase catalyses the dephosphorylation of sedoheptulose-1,7-bisphosphate, the reaction nested at the branch point between the regenerative phase of the C_3_ cycle and sucrose or starch biosynthesis. Due to this unique position, SBPase is one of the critical enzymes controlling the carbon flow in plants (Raines et al., 2000).

A major limitation of the C_3_ cycle is the enzyme ribulose-1,5-bisphosphate carboxylase/oxygenase (Rubisco) catalysing the fixation of CO_2_ into ribulose-1,5-bisphosphate (RuBP) producing glycerate-3-phosphate, a 3-C compound giving the name ‘C_3_ species’ to those that use this cycle exclusively. However, Rubisco also catalyses an oxygenase reaction that competes with CO_2_ fixation resulting in reductions in yield of over 25% (Walker et al., 2016). To get around this problem, ~ 4 % of plant species have evolved a biochemical CO_2_ concentrating mechanism, called the C_4_ pathway, that operates in addition to the C_3_ cycle and involves two functionally distinct cell types. In the C_4_ pathway, atmospheric CO_2_ diffuses into the leaf mesophyll cells where it is converted to HCO_3_^−^ by carbonic anhydrase (CA) which is then fixed by phosphoenolpyruvate (PEP) carboxylase (PEPC) to produce C_4_ acids (giving the name to this pathway). These C_4_ acids diffuse into the bundle sheath cells where they are decarboxylated, thereby elevating the CO_2_ partial pressure (*p*CO_2_) where Rubisco is located, allowing Rubisco to operate close to its maximal rate (Hatch, 1987). C_4_ crops are high yielding and are characterised by high photosynthetic rates, high nitrogen and water use efficiency when compared to plants using only the C_3_ cycle. This has stimulated considerable interest in the C_4_ photosynthetic pathway (Mitchell and Sheehy, 2006), and a range of strategies to manipulate and enhance C_4_ photosynthesis are also being considered (von Caemmerer and Furbank, 2016). The rate of Rubisco catalysis and the regeneration of PEP and RuBP together with the electron transport capacity are all possible limiting factors of C_4_ photosynthesis under high *p*CO_2_ and high irradiance conditions (von Caemmerer and Furbank, 2016).

To explore these limitations, a new model species, *Setaria viridis* (green foxtail millet), has emerged to enable the study of C_4_ plant biology. Like the major C_4_ crops maize (*Zea mays*) and sorghum, *S. viridis* belongs to the NADP-ME decarboxylation C_4_ subtype and is readily transformable opening opportunities to investigate the possibility of enhancing C_4_ photosynthesis (Brutnell et al., 2010). Overexpression of the Rieske FeS protein in *S. viridis* has been shown to enhance C_4_ photosynthesis (Ermakova et al., 2019) and the joint overexpression of Rubisco subunits with the RUBISCO ASSEMBLY FACTOR 1 in *Z. mays* has increased Rubisco protein content and photosynthetic rate (Salesse-Smith et al., 2018). Given that the C_3_ cycle plays an equally important role in C_4_ plants and the success reported in enhancing C_3_ photosynthesis by increasing SBPase content, here we investigated whether overexpression of SBPase could also enhance C_4_ photosynthesis. To test this hypothesis, we produced and analysed *S. viridis* plants expressing *Brachypodium distachyon SBPase* using a bundle sheath cell-preferential promoter. Our results showed that SBPase content does not limit C_4_ photosynthetic flux under any of the environmental conditions tested.

## Materials and Methods

### Generating transgenic plants

The coding sequence of *Brachypodium distachyon SBPase* (Bradi2g55150, https://phytozome-next.jgi.doe.gov) was codon-optimised for the Golden Gate cloning system (Engler et al., 2014) and assembled with the bundle sheath cell-preferential *Flaveria trinervia GLDP* (glycine decarboxylase P-protein) promoter (Engelmann et al., 2008, Gupta et al., 2020) and the bacterial *NOS* (nopaline synthase) terminator. The resulting expression module was cloned into the second slot of a plant binary vector pAGM4723. The first slot was occupied by the *hpt* (hygromycin phosphotransferase) gene driven by the *Oryza sativa Actin1* promoter. The construct was verified by sequencing and transformed into *Setaria viridis* cv. MEO V34-1 using *Agrobacterium tumefaciens* strain AGL1 according to the protocol described in detail in Osborn et al. (2016). T_0_ plants resistant to hygromycin were transferred to soil and tested for SBPase abundance by immunoblotting and *hpt* copy number by droplet digital PCR (iDNA genetics, Norwich, UK). Wild type (WT) plants were used as control in all experiments.

### Plant growth conditions

Seeds were surface-sterilized and germinated on rooting medium containing 2.15 g L^−1^ Murashige and Skoog salts, 10 ml L^−1^ 100x Murashige and Skoog vitamins stock, 30 g L^−1^ sucrose, 7 g L^−1^ Phytoblend, 20 mg L^−1^ hygromycin (no hygromycin for WT plants), pH 5.7. Seedlings that developed secondary roots were transferred to 1 L pots with the commercial potting mix (Debco, Tyabb, Australia) layered on top with 2 cm of the seed raising mix (Debco) both containing 1 g L^−1^ Osmocote (Scotts, Bella Vista, Australia). Plants were grown in controlled environment chambers with ambient CO_2_, 16 h photoperiod, 28 °C day, 22 °C night and 60 % humidity. Light at the intensity of 300 μmol m^−2^ s^−1^ was supplied by 1000 W red sunrise 3200K lamps (Sunmaster Growlamps, Solon, OH). Youngest fully expanded leaves of 3 weeks-old plants were used in all experiments. Photosynthetic and physiological parameters of leaves were measured with the MultispeQ using ‘Photosynthesis RIDES’ protocol at ambient conditions in the growth chamber (Kuhlgert et al., 2016). The results were analysed using the PhotosynQ platform (https://photosynq.com).

### Immunoblotting

Leaf discs of the same area were collected and immediately frozen in liquid N_2_. Protein samples were isolated from leaf discs as described in Ermakova et al. (2019). Proteins were separated by SDS-PAGE, transferred to a nitrocellulose membrane and probed with antibodies against SBPase (AS152873, Agrisera, Vännäs, Sweden), RBCL (Martin-Avila et al., 2020) and PEPC (Karki et al., 2020). Quantification of immunoblots was performed with the Image Lab software (Biorad, Hercules, CA).

### Bundle sheath isolation and qPCR

BS strands were isolated following the procedure of Ghannoum et al. (2005) as described in detail in Ermakova et al. (2021a). RNA was isolated from leaves and BS strands, ground in liquid N_2_, using the RNeasy Plant Mini Kit (Qiagen, Venlo, The Netherlands). DNA was removed from the samples using the Ambion TURBO DNA free kit (Thermo Fisher Scientific, Tewksbury, MA). cDNA was synthesised and analysed by qPCR as described in (add ref Ermakova 2019). Relative fold change was calculated by the ΔΔC_t_ method using the geometric mean of the C_t_ values for three reference genes described in Osborn et al. (2016). Primers to distinguish between *S. viridis* and *B. distachyon SBPase* transcripts were designed using Primer3 in Geneious R9.1.1 (https://www.geneious.com).

### Gas Exchange

Gas exchange analysis was performed using a LI-6800 (LI-COR Biosciences, Lincoln, NE). First, leaves were equilibrated at 1500 μmol m^−2^ s^−1^ (90 % red / 10 % blue actinic light), 400 ppm CO_2_ in the reference side, leaf temperature 28 °C, 60% humidity and flow rate of 500 μmol s^−1^, and then light or CO_2_ response curves of CO_2_ assimilation were recorded. For the light response curves, a stepwise increase of irradiance from 0 to 3000 μmol m^−2^ s^−1^ was imposed at 2-min intervals. For the CO_2_ response curves, a stepwise increase of CO_2_ partial pressures from 0 to 1600 ppm was imposed at 3-min intervals. To record CO_2_ response curves at different temperatures, plants were kept in growth cabinets set to 15 °C or 35 °C, and leaves were equilibrated at a corresponding leaf temperature for 20 min before the measurement.

### Statistical analysis

The relationship between mean values of transgenic and WT plants was tested using two-tailed, heteroscedastic Student’s *t*-test.

## Results

Six *S. viridis* plants resistant to hygromycin were regenerated after the transformation with the construct for SBPase overexpression. The *hpt* copy number identified by the digital PCR indicated that T_0_ plants contained one to three *B. distachyon SBPase* (*BdSBPase*) copies (Fig. 1a). Immunodetection of SBPase and PEPC suggested increased SBPase abundance, relative to PEPC, in multiple T_0_ plants when compared to WT. Plants 2, 3 and 5 were selected and the progeny of these lines was further analysed. To verify bundle sheath cell-preferential expression of *BdSBPase*, total RNA was isolated from leaves and bundle sheath strands of WT plants and homozygous T_1_ plants of line 3. Transcript abundance of *BdSBPase* and the native *SBPase* (*SvSBPase*) in the bundle sheaths exceeded the levels detected from whole leaves indicating that both genes were preferentially expressed in bundle sheath cells (Fig. 1b).

**Figure 1.**
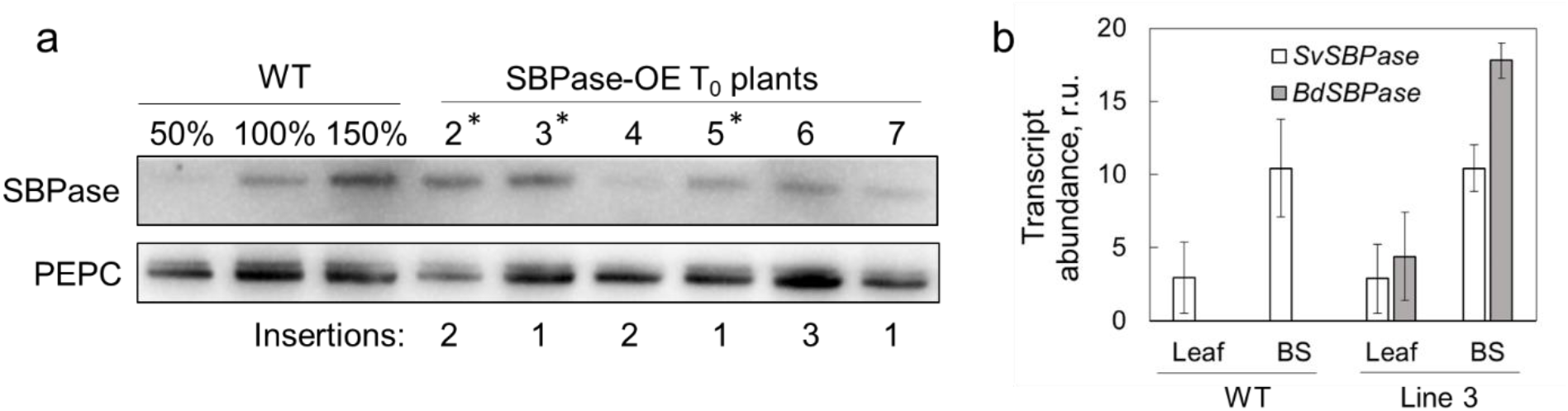
Selection of *S. viridis* plants overexpressing SBPase. **a**. Immunoblots of SBPase and PEPC in WT *S. viridis* and 6 T_0_ plants transformed with *SBPase* from *Brachypodium distachion* under the control of the bundle sheath cell-preferential *GLDP* promoter. The *htp* copy numbers estimated by digital PCR and suggesting the *BdSBPase* insertion numbers are also shown. Asterisks indicate the plants which progenies were used in further experiments. **b**. Transcript abundance of *S. viridis SBPase* (*SvSBPase*) and *B. distachion SBPase* (*BdSBPase*) in whole leaf tissue and isolated bundle sheath strands (BS) shows bundle sheath-preferential localisation of the native gene and transgene transcripts. Mean ± SE, *n* = 3 biological replicates.

We studied the impact of increased SBPase abundance on C_4_ photosynthesis first by analysing T_1_ plants of lines 2 and 3 for *hpt* insertion number and SBPase and Rubisco large subunit (RBCL) content. Relative abundance of SBPase, quantified from the immunoblots (Fig. 2a), showed strong positive correlation with the insertion number in T_1_ plants and, thus, with the copy number of *BdSBPase* (Fig. 2b). Homozygous plants of line 2, containing four copies of *BdSBPase*, showed the highest SBPase levels (about four times of WT). Relative content of RBCL showed no correlation with the insertion number (Fig. 2c) indicating that neither increased SBPase abundance or insertion positions had an impact on Rubisco abundance in transgenic plants. In addition, neither relative Chl abundance or leaf thickness was affected in plants with increased SBPase content, compared to WT (Table 1).

**Figure 2.**
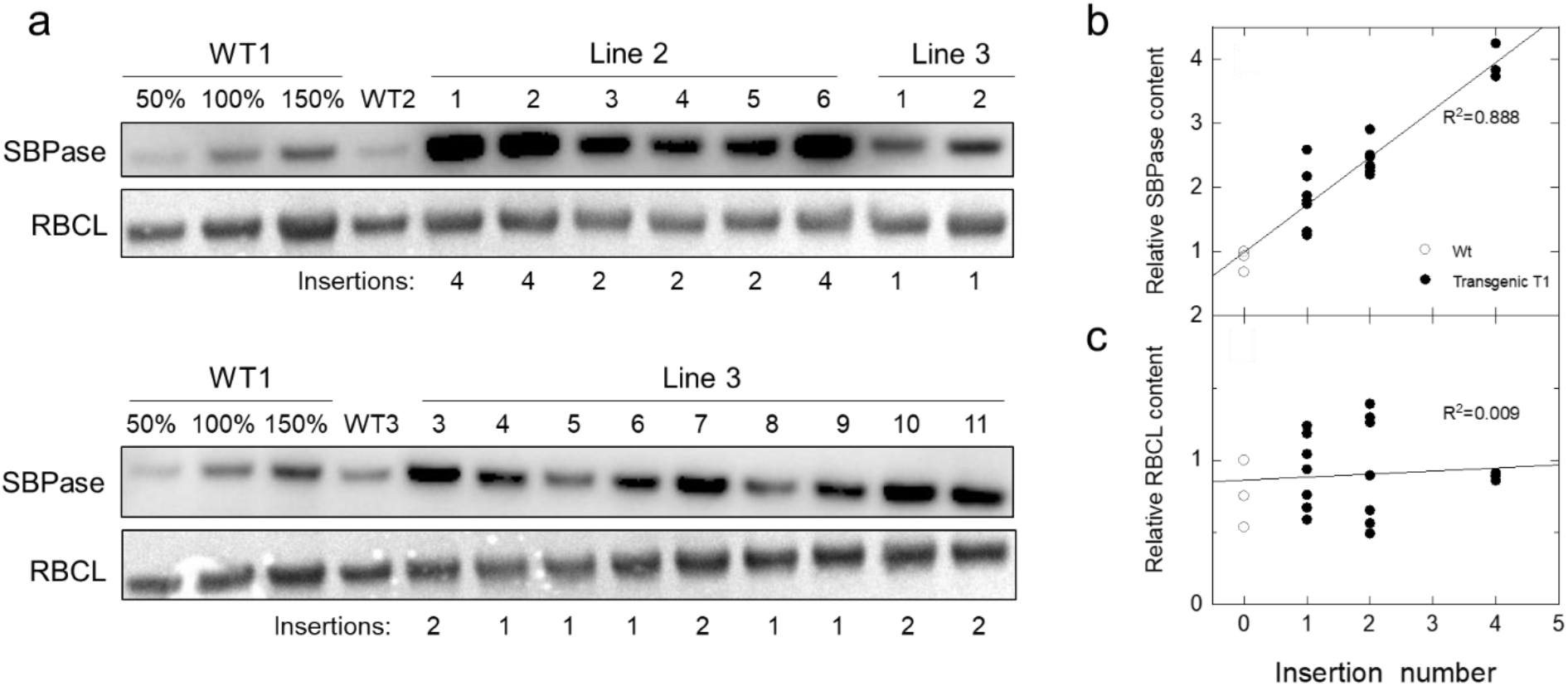
Analysis of *S. viridis* plants overexpressing SBPase. **a**. Immunodetection of SBPase and RBCL in protein samples isolated from leaves of WT plants and the T_1_ progeny of transgenic lines 2 and 3. Samples were loaded on leaf area basis, and the titration series of the WT1 sample was used for relative quantification. Insertion number indicates *hpt* copy number estimated by digital PCR. **b** and **c**. Relative SBPase and RBCL content as a function of insertion number (data taken from a).

**Table 1.** Photosynthetic and physiological parameters measured on leaves of wild type (WT) *S. viridis* and two transgenic lines overexpressing SBPase at growth light intensity. PhiPSII, the effective quantum yield of Photosystem II; PhiNPQ, the yield of non-photochemical quenching; PhiNO, the yield of non-regulated non-photochemical reactions in PSII; g_H+_, proton conductivity of the thylakoid membrane. Mean ± SE, n = 6-7 biological replicates. No statistically significant differences were found between transgenic and WT plants (*P* < 0.05).

| Parameter | WT | Line 2 | Line 3 |
| --- | --- | --- | --- |
| Relative Chl (SPAD) | 43.57± 0.83 | 42.02± 1.18 | 43.00± 0.95 |
| Leaf thickness, mm | 0.59± 0.08 | 0.64± 0.07 | 0.58± 0.09 |
| PhiPSII | 0.62± 0.01 | 0.62± 0.01 | 0.63± 0.02 |
| PhiNO | 0.21± 0.01 | 0.20± 0.01 | 0.20± 0.01 |
| PhiNPQ | 0.17± 0.01 | 0.18±0.01 | 0.17± 0.01 |
| $g_{H^+}$ | 240.4± 18.2 | 239.4± 7.2 | 255.1± 15.2 |

Next, we studied photosynthetic properties of *S. viridis* plants with increased SBPase content. Figure 3 shows that CO_2_ assimilation rates measured from WT and T_1_ plants of lines 2 and 3 at ambient CO_2_ partial pressure were not affected by SBPase content, quantified from the immunoblots (Fig. 2a). Moreover, no difference in CO_2_ assimilation rates was detected between WT and transgenic plants overexpressing SBPase at different intercellular CO_2_ partial pressures or irradiances (Fig. 4). Electron transport parameters measured at growth light intensity indicated that plants with increased SBPase content had WT-like activity of Photosystem II, since no changes in partitioning of the absorbed light between photochemical (PhiPSII) and non-photochemical (PhiNPQ and PhiNO) reactions within Photosystem II were detected (Table 1). Activity of the chloroplast ATP synthase, estimated as the proton conductivity of thylakoid membrane (g_H+_), did not differ between the genotypes either (Table 1).

**Figure 3.**
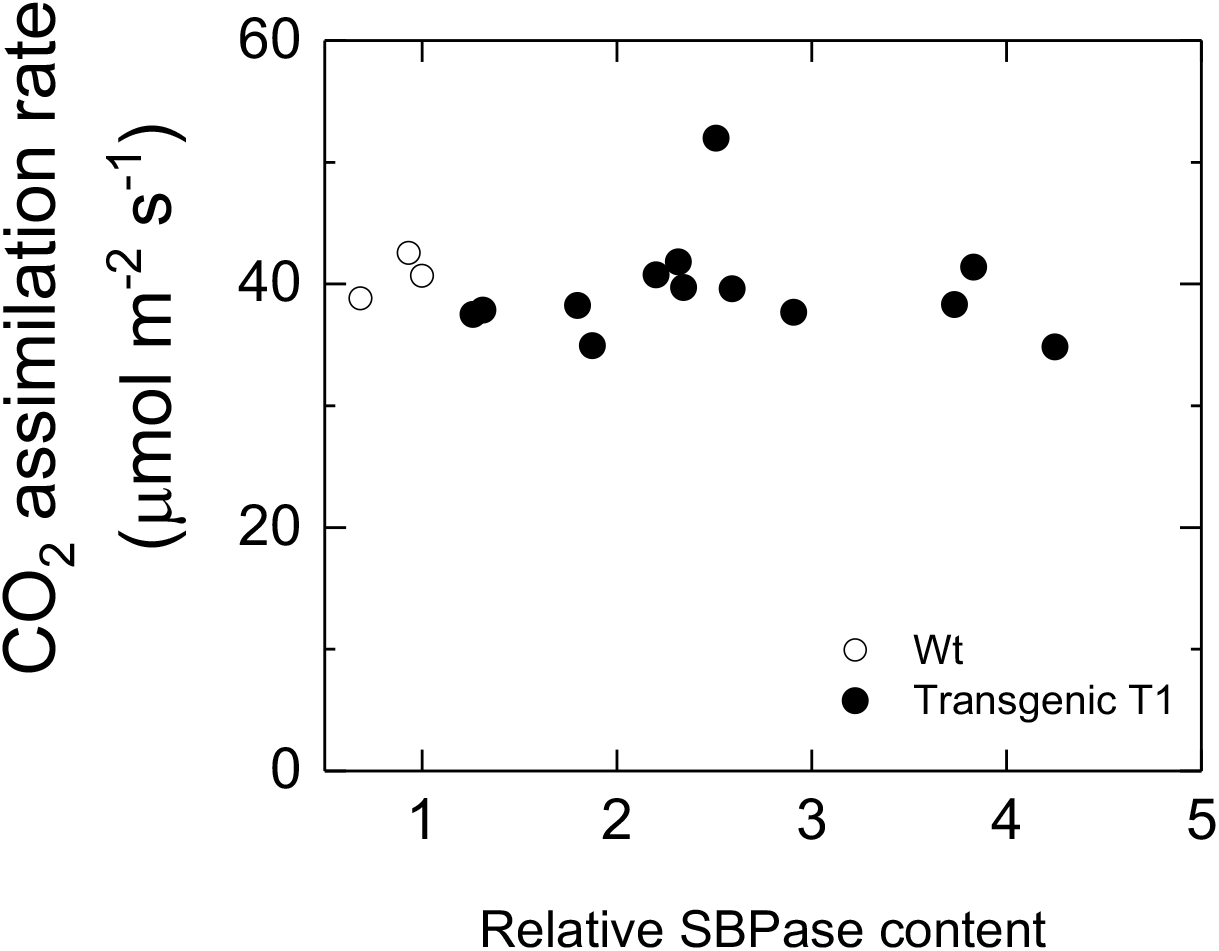
Saturating CO_2_ asimilation rate as a function of relative SBPase content in leaves of WT *S. viridis* and T_1_ progeny of lines 2 and 3 overexpressing SBPase. Measurements were made at an ambient CO_2_ partial pressure of 400 μbar, an irradiance of 1500 μmol m^−2^ s^−1^ and a leaf temperature of 28 °C.

**Figure 4.**
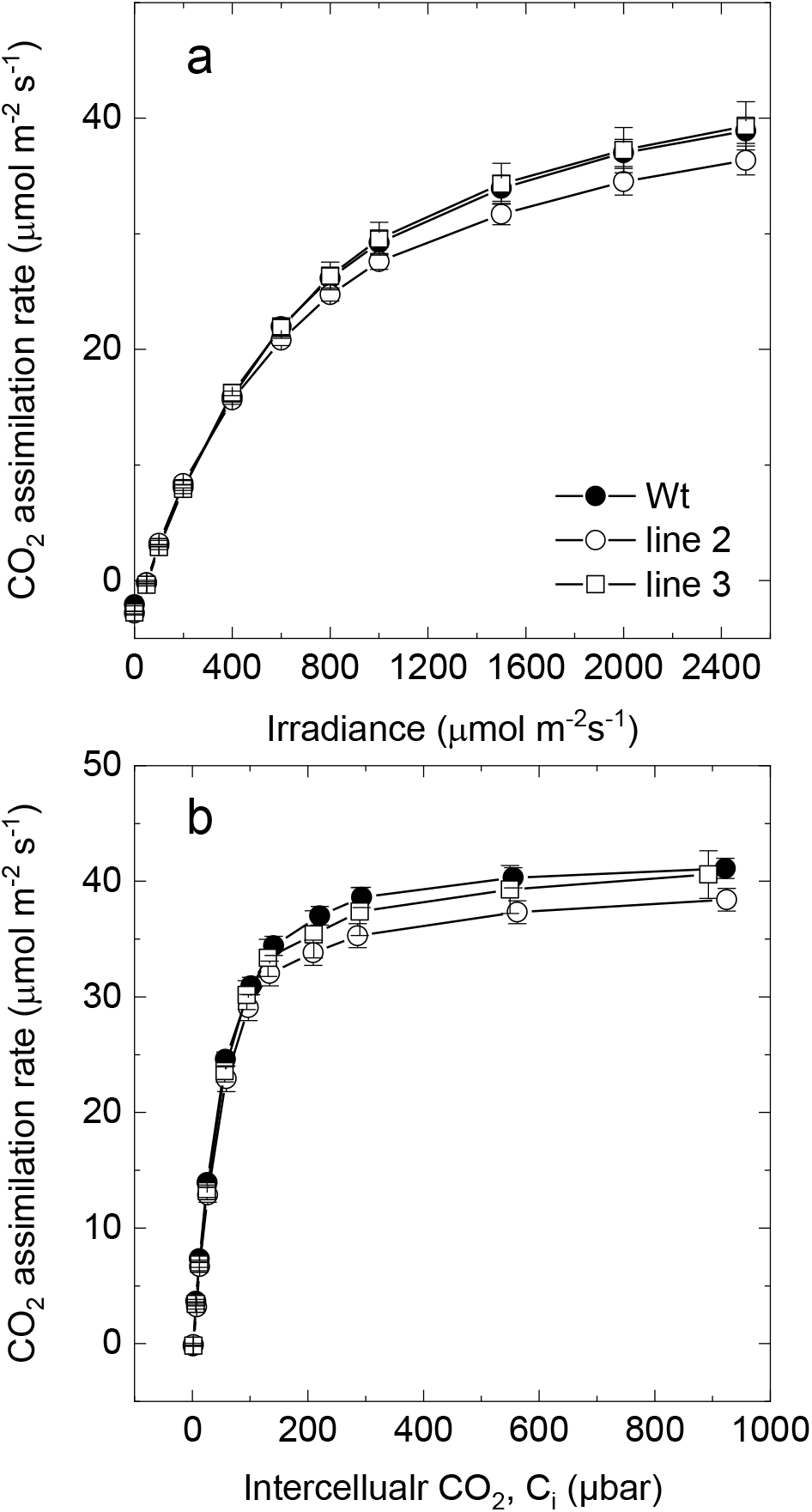
Gas exchange properties of WT *S. viridis* and transgenic plants overexpressing SBPase. **a**. CO_2_ assimilation rate as a function of intercellular CO_2_ partial pressure. Measurements were made at an irradiance of 1500 μmol m^−2^ s^−1^ and a leaf temperature of 28 °C. **b.** CO_2_ assimilation rate as a function of irradiance. Measurements were made at an ambient CO_2_ partial pressure of 400 μbar and a leaf temperature of 28 °C. Measurments were made on the T_1_ progeny of lines 2 and 3. Average relative abundance of SBPase per leaf area, calculated from immunoblots on Fig. 2, was significantly increased in line 2 (3.2 times, *P* = 0.002) and in line 3 (2.0 times, *P* = 0.001), relative to WT. Mean ± SE, *n* = 3 biological replicates. No significant differences were found (*P* < 0.05).

We also tested CO_2_ assimilation rates at different temperature in homozygous T_2_ plants of lines 3 and 5 with increased SBPase abundance confirmed by immunoblotting (Fig. S1). For this, CO_2_ response curves of assimilation were measured on leaves acclimated to 35 °C and at 15 °C (Fig. 5). No difference in CO_2_ assimilation was detected between WT and transgenic plants overexpressing SBPase at 35 °C. At 15 °C, both transgenic lines showed WT-like rates of assimilation, except for the plants of line 5 having significantly increased CO_2_ assimilation rate of 6.80 ± 0.04 μmol m^−2^ s^−1^ at the intercellular CO_2_ partial pressure of about 13 μbar, compared to the WT rate of 5.14 ± 0.35 μmol m^−2^ s^−1^ (*P* = 0.040, *t*-test).

**Figure 5.**
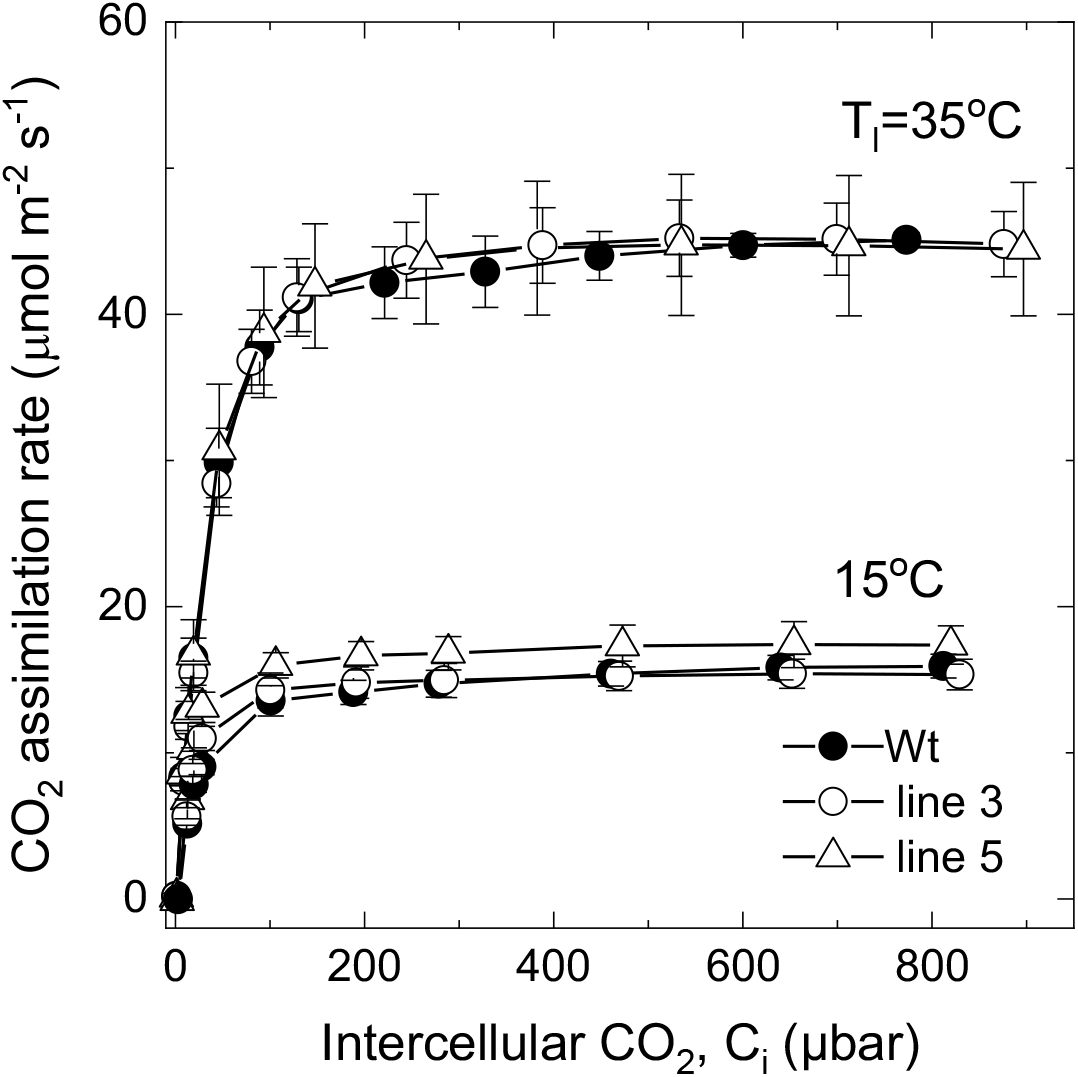
CO_2_ assimilation rate as a function of intercellular CO_2_ partial pressure at different temperatures in WT *S. viridis* and transgenic plants over-expressing SBPase. Measurements were made at an irradiance of 1500 μmol m^−2^ s^−1^ and a leaf temperature of 35 °C or 15 °C. Measurments were made on the T_2_ progeny of lines 3 and 5 and wild type. Average relative abundance of SBPase per leaf area, calculated from immunoblots on Fig. S1, was increased 2 times in line 3 and 1.5 times in line 5, relative to WT. Mean ± SE, *n* = 3 biological replicates. Details of statistical analysis are provided in the results.

## Discussion

Overexpression of SBPase has led to increases in photosynthetic rates and increased biomass in several C_3_ species (Rosenthal et al., 2011, Lefebvre et al., 2005, Driever et al., 2017, Feng et al., 2007, Ding et al., 2016). In the C_4_ photosynthetic system, SBPase and the C_3_ cycle are located in the bundle sheath cells, and therefore the C_3_ cycle in C_4_ photosynthesis is poised differently to the C_3_ cycle in C_3_ species. Because the C_3_ cycle operates at high CO_2_ partial pressure in the bundle sheath, C_4_ species have characteristically less Rubisco protein then C_3_ species (Sage and Pearcy, 1987, von Caemmerer and Furbank, 2016), however to achieve high photosynthetic rates SBPase levels will need to be similar to that in C_3_ species.

Here we expressed *BdSBPase* from a bundle sheath cell-preferential promoter to test SBPase overexpression in a C_4_ photosynthetic system. We observed increased SBPase content in T_0_ plants and selected 3 T_1_ lines for our investigations (Fig. 1). Transgenic plants had between 1.5 to 3.2 times the amount of SBPase, relative to WT, as judged from the immunoblots (Fig. 2a, Fig. S1). Despite these significant increases in protein content, we observed no increase in photosynthetic rates under a range of environmental conditions including different irradiances, *p*CO_2_ and temperatures. The C_4_ photosynthetic model suggests that SBPase limitation should be apparent at *p*CO_2_ above ambient where SBPase content may well be co-limiting with electron transport capacity, Rubisco activity and PEP regeneration (von Caemmerer, 2021, von Caemmerer and Furbank, 1999).

*S. viridis* uses NADPH-dependent malic enzyme (NADP-ME) decarboxylation system in the bundle sheath chloroplast (Fig. 6) and, similar to most of NADP-ME species, *S. viridis* has low PSII activity and linear electron transport rate in bundle sheath cells (Ermakova et al., 2021a). This prompts export of part of the 3-phosphoglycerate (3-PGA) pool to the mesophyll for conversion to triose phosphate, which then diffuses back to the bundle sheath, known as the triose phosphate shuttle (von Caemmerer and Furbank, 2016 and references therein) (Fig. 6). To support this movement of triose phosphate and 3-PGA, diffusion gradients between the mesophyll cells and the bundle sheath must be built up and maintained (Furbank and Kelly, 2021 and references therein). Moreover, reactions of sucrose and starch synthesis are distributed between the different cell types in the C_4_ system with sucrose being made mostly in the mesophyll cells and starch - in the bundle sheaths (Furbank et al., 1985, Lunn and Furbank, 1997, Furbank and Kelly, 2021). The combination of the triose phosphate shuttle and the cellular localisation of sucrose and starch biosynthesis may mean that regulation of the regeneration of RuBP in C_4_ bundle sheath chloroplasts is somewhat different to that in C_3_ chloroplasts where sucrose and starch synthesis and 3-PGA reduction are all occurring in a single cell type. It has been proposed that both SBPase and FBPase play key roles in determining the fate of triose phosphate in the C_3_ cycle, *i.e*., whether it is recycled to regenerate RuBP or used to make sucrose and starch (Raines et al., 2000 and references therein). This regulation is important as the metabolite pools within the cycle need to be preserved and flux maintained while carbon is removed for storage. Because of the, this regulation is controlled not only by the activities of these biphosphatases but by a complex balance of orthophosphate consumption in photophosphorylation and its resupply from P_i_ release in sucrose biosynthesis and the activity of ADPG-pyrophosphorylase (Furbank and Kelly, 2021). Since in the C_4_ bundle sheath chloroplast, triose phosphate is imported in exchange for 3-PGA export, the P_i_ recycling process is by necessity different to the C_3_ case where triose phosphate is exchanged for P_i_. This division of metabolism between the cell types and the higher flexibility of carbon flow in C_4_ plants might reduce the capacity to regulate carbon flux by the abundance of SBPase.

**Figure 6.**
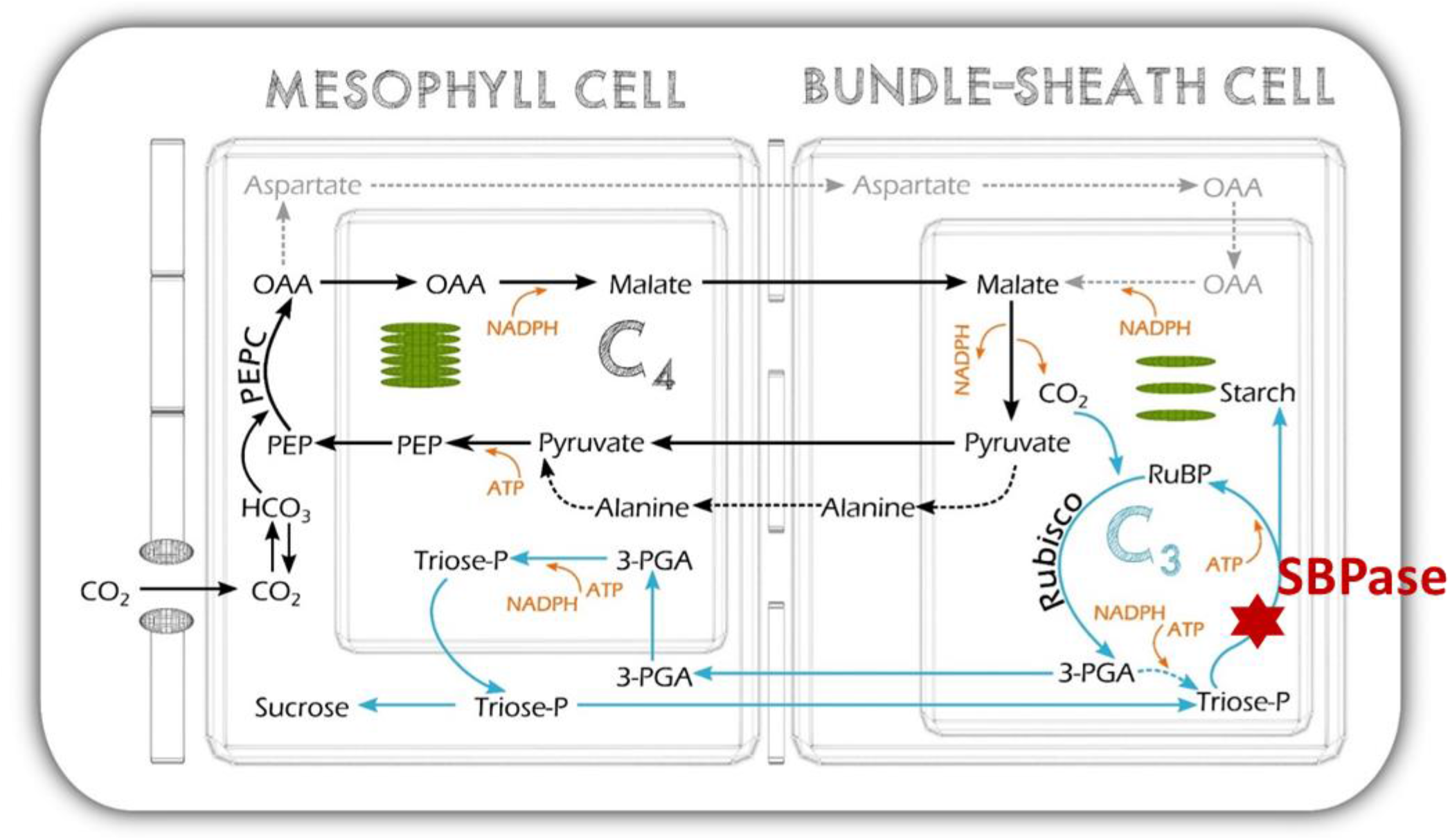
Schematic of the NADP-ME C_4_ photosynthetic pathway of *S. viridis* showing the location of SBPase in the pathway.

Little is known about potential differences in regulation of SBPase between C_3_ and C_4_ plants. There is evidence that the kinetic properties of enzymes such as cytosolic FBPase are quite different in C_4_ plants to support the cellular gradients required for fluxes of metabolites (Furbank and Kelly, 2021). Wheat SBPase was shown to be regulated by pH and Mg^2+^ concentrations consistent with its activity being stimulated under light (Woodrow et al., 1984). Moreover, like some other enzymes of the C_3_ cycle, SBPase is activated by the thioredoxin system via a light-dependent reduction of the disulfide bond (Breazeale et al., 1978, Dunford et al., 1998). Since we were not able to measure *in vitro* activity of SBPase, there is a possibility that SBPase from *B. distachyon*, a C_3_ plant, was inactive when expressed in the bundle sheath cells of C_4_ plant. However, the amino acid sequences of SBPase from *S. viridis* and *B. distachyon* are 92.4 % identical, and all cysteine residues are conserved (Fig. S2). Moreover, SBPase from *Z*. *mays* could be activated by thioredoxin *f* from spinach suggesting a cross-reactivity between enzymes and thioredoxins from different species (Nishizawa and Buchanan, 1981). Nevertheless, although a positive correlation was typically observed between the active form and total enzyme abundance in C_3_ plants overexpressing SBPase (Driever et al., 2017), there is a possibility that the ‘extra’ SBPase is not activated in *S. viridis* due to a limited availability of reducing power to the thioredoxin system in C_4_ bundle sheath cells. In that case, C_4_ photosynthesis would not be limited by SBPase directly, but rather by electron transport, which has been previously confirmed (Ermakova et al., 2019).

## Conclusion

Under the range of conditions tested in this study, increasing SBPase levels did not increase photosynthetic flux in the C_4_ grass *Setaria viridis*, in contrast to observations of SBPase overexpression in C_3_ plants. We propose that this is because of (i) the triose phosphate shuttle in C_4_ plants, where part of the 3-PGA produced by Rubisco is reduced in the mesophyll chloroplasts and returned to the bundle sheath chloroplasts, and (ii) the cellular specialisation of starch and sucrose biosynthesis in C_4_ leaves where these processes are spatially separated. These unique aspects of the C_4_ photosynthetic pathway are likely to result in a different distribution of control over regeneration of RuBP and the coordination of RuBP production and sugar phosphate utilisation in starch and sucrose synthesis. At present, the degree of sophistication of the C_4_ photosynthetic model is insufficient to accommodate these finely tuned control mechanisms.

## Acknowledgements

We thank Xueqin Wang for help with *S. viridis* transformation, Zac Taylor and Ayla Manwaring for immunoblotting and Emily Watson for gas exchange measurements. This work was supported by the Australian Research Council Centre of Excellence in Translational Photosynthesis (CE140100015).

## Author Contributions

SvC, CR, RF and ME designed the research; ME and PL performed research; ME and SvC analysed data; SvC, ME, CR and RF wrote the paper.

## Conflict of Interest

Authors declare no conflict of interest.

